# BEE : a web service for biomedical entity exploration

**DOI:** 10.1101/2020.01.03.893594

**Authors:** Jin-uk Jung, Jin-Muk Lim, Hyunwhan Joe, Hong-Gee Kim

## Abstract

Recently there has been a trend in bioinformatics to produce and manage large quantities of data to better explain complex life phenomena through relationship and interactions among biomedical entities. This increase in data leads to a need for more efficient management and searching capabilities. As a result, Semantic Web technologies have been applied to biomedical data. To use these technologies, users have to learn a query language such as SPARQL in order to ask complex queries such as ‘What are the drugs associated with the disease breast carcinoma and Osteoporosis but not the gene ESR1’. BEE was developed to overcome the limitations and difficulties of learning such query languages. Our proposed system provides an intuitive and effective query interface based on natural language. Our system is a heterogeneous biomedical entity query system based on pathway, drug, microRNA, disease and gene datasets from DGIdb, Tarbase, Human Phenotype Ontology and Reactome, Gene Ontology, KEGG gene set of MSigDB. User queries can be joined with union, intersection and negation operators. The system also allows for selected results to be saved and later combined with newly created queries. To the best of our knowledge, BEE is the first system that supports condition search based on the relationship of heterogeneous biomedical entities and is expected to be used in various fields of bioinformatics such as in drug repositioning candidate selection as well as simple knowledge search.

## Introduction

In the last decades, bioinformaticians have been trying to discover the relationships between genes and other biomedical entities. As a result, the amount of data on gene associations with biomedical entities such as diseases, drugs and pathways have exploded. Consequently, this growth in data has lead to an increasing need for an efficient search tool for the relationship between more complex biomedical entities.

BEE is a web service that was developed for the needs of these users. The system is an entity search system that can answer simple questions such as ‘what are the pathways containing gene A and gene B’, to more complex questions such as ‘what are the drugs associated with disease *s*_1_ and *s*_2_ without gene *g*_1_ targeted drugs’, which requires knowledge between heterologous biomedical entities. Our primary goal is to develop an easy-to-create query system that can get results for complex queries. To search with multi-queries in existing systems, the user needs to learn a query language such as SPARQL or adapt to the use of the complex interface provided by the system. It results in a deterioration of user accessibility. The design principle of BEE is focused on intuitiveness and simplicity. BEE’s straightforward interface minimizes the level of questions and input parameters, reducing the difficulty for new users. In addition, the system can be easily expanded by minimizing the difficulty of adding and modifying a new biomedical entity with a simple data structure.

## MATERIALS AND METHODS: Inside of BEE

### A. Design principle

The development focus was to minimize the difficulty of query generation for users, which was the limitation of existing search systems. Therefore, development proceeded by setting up the interface to be as simple and intuitive as possible.

Subsequent considerations were focused on minimizing the query step and reducing the required parameters. Also, for intuitive use, the parameter input follows the order of words in natural language. This minimizes the learning curve of how to use the system and allows the user to use the application immediately without any difficulties. The system provides features that save/load the search history, and supports exporting search results into various formats such as clipboard, CSV, and Excel. In addition, BEE has a flexible schema that allows developers to add existing and new biomedical entity types, while ensuring that there is no impact of change to the existing system.

### B. Design principle

There are five entity types used in BEE: gene, disease, drug, pathway and microRNA. Drug-gene interaction data were extracted from DGIdb(1). DGIdb is a dataset that integrates human genes and drug-related data related to diseases based on 13 major sources. It provides more than 14,000 drug-gene interaction data between more than 2,600 genes and 6,300 drugs. It was last updated on 2018-01-25. The gene symbol, Entrez ID, drug name and ChEmBL ID included in the schema were extracted and loaded into the BEE database. Second, disease-gene association is based on data provided by the Human Phenotype Ontology(HPO)(2). HPO provides a standard vocabulary of phenotypic features of human genetic or other diseases. HPO data is updated once a month, and the preprocessing module in our system extracts the entrez id, gene symbol, HPO term, and HPO term ID data from the files provided by HPO and stores them in our database. Third, microRNA data is based on TarBase(3). TarBase provides sequence data, target gene information and annotations on a web service, and integrates information on cell-type specific miRNA-gene regulation information. Our preprocessing module extracted the gene symbol and miRNA information from the data provided by the source, and filtered it by the species Homo sapiens. TarBase was last updated on 2018-3-12. Fourth, the pathway-gene association data was collected from the Gene Ontology(Biological process)(2) (4), Reactome(5), and KEGG(6) Gene set data is provided by the Molecular Signatures Database(MSigDB)(7). MSigDB provides only gene information for each pathway and removes the context. Our system imported data which was released in MSigDB 7.0. Finally, gene data was composed by integrating all gene symbols linked to the above disease, drug, pathway, and microRNA. The BEE database consists of 25,033 genes, 20,370 diseases, 5256 pathways, 12785 drugs, and 1034 microRNAs.

### C. System model

A Model of BEE *B* = (*G, E, R*_*E*−*G*_). *G* is a set of gene {*g*_1_, *…, g*_*v*_}. *E* is a set of entity types {*D, S, P, M*}. It includes a set of drugs *D*{*d*_1_, *…, d*_*w*_}, a set of diseases *S*{*s*_1_, *…, s*_*x*_}, a set of pathways *P* {*p*_1_, *…, p*_*y*_}, and a set of microRNAs *M* {*m*_1_, *…, m*_*z*_}. And *R*_*E*−*G*_ is a set of relations between *e*_*i*_ ∈ *E* and *g*_*i*_ ∈ *G* that is *R*_*E*−*G*_ ⊆ *E* × *G*. For example, as shown in Figure 1, given *d*_*w*_ ∈ *D* and 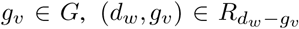 means “drug *d*_*w*_ target to gene *g*_*v*_”. i.e. *R*_*D*−*G*_ = {(*g*_1_, *d*_1_), (*g*_1_, *d*_2_)}, *R*_*S*−*G*_ = {(*g*_1_, *s*_1_), (*g*_2_, *s*_2_), (*g*_3_, *s*_2_), (*g*_4_, *s*_2_)}.

**Fig. 1.**
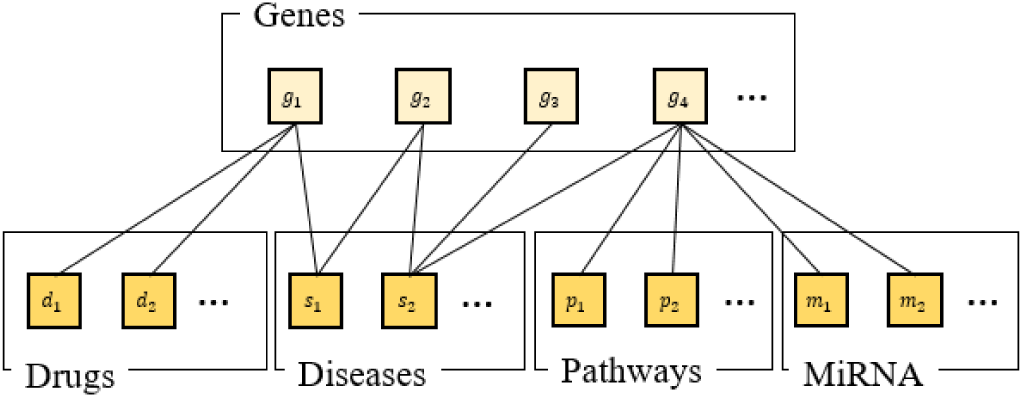
Entity association in BEE.

And also, the system model has three functions, *gen, rgen, ext*. First, *gen*, given an entity *e*_*i*_ ∈ *E*, 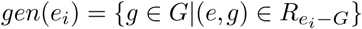. The function takes an element of entity as parameter, returns the gene set associated with the input entity. E.g. *gen*({*s*_1_}) = {*g*_1_, *g*_2_}. Second, given a gene *g*_*i*_ ∈*G, rgen*(*g*_*i*_) = {{*e*} ⊆ *|*(*e, g*) ∈ *R*_*E*−*G*_}. The function *rgen* takes an element of gene entity as a parameter and returns all entities associated with the input gene. E.g. *rgen*({*g*_1_}) = {*d*_1_, *d*_2_, *s*_1_}, *rgen*({*g*_2_}) = {*s*_1_, *s*_2_}. Third, the *ext* function extracts and returns a specific type of entity from the set of input entities. Given a set of entities *e*_*i*_ ∈ *E* and entity type *T, ext*(*T, E*), for example *ext*(*T*_*D*_, {*s*_1_, *s*_2_, *d*_1_, *p*_1_}) = {*d*_1_}. In essence, the system model of BEE is a web service that provides elements to users by sequentially processing the three operators as *ext*(*T*_*D*_, *rgen*(*gen*({*s*_1_}))).

### D. Web server development

BEE is developed under the Laravel framework and works with MySQL databases. Laravel is a PHP web framework based on the MVC pattern. It supports the flexibility of database changes, high security, and a lightweight, separated template engine. The user interface was based on Bootstrap (https://getbootstrap.com/). Bootstrap provides various web page layouts, buttons, input windows, and icons in CSS and Javascript for rapid development, and cross-browsing and converting a single web page into an optimized layout for desktop or mobile devices. These features help to shorten front-end development. The entity graph that appears in the search results uses force-graph (https://github.com/vasturiano/force-graph), and dynamic tables are implemented using DataTables (https://datatables.net/).

## Results

### E. BEE web server

The system is based on the gene, drug, disease, pathway, and microRNA database loaded by the preprocessing module, and the loaded data continues to grow as the data sources in each silo update.

On the first screen of BEE, the user can enter a query. As a result, the system provides network visualization information and query results between entities according to the input query, and the query result can be saved and used as part of the query chain.

### F. Query interface

The system provides an interface for getting answers by combining two or more query sets and their operators. There are three elements that make up a query set: Answer entity type, Question entity type and Question entity name. As shown in Table 1, in query set 1, for the question “What are the drugs associated with Breast carcinoma”, the Answer entity type is ‘Drug’. ‘Disease’ is the question entity type and ‘Breast carcinoma’ is the Question entity name. There are also three query operators which are union, intersection and negation. For example, the question ‘What are the drugs associated with disease Breast carcinoma and Osteoporosis but not gene ESR1’ can be obtained by setting “query set1 ∩ query set2 ¬ query set3” in the search operators section.

**Table 1.** User query set example

|  |  |  |  |  |
| --- | --- | --- | --- | --- |
| 3*<br>Query<br>set 1 | Parameter | Answer entity type | Question entity type | Question entity name |
|  | Input value | Drug | Disease | Breast carcinoma |
|  | Natural Language | What are drugs that associate with Breast carcinoma? |  |  |
| 3*<br>Query<br>set 2 | Parameter | Answer entity type | Question entity type | Question entity name |
|  | Input value | Drug | Disease | Osteoporosis |
|  | Natural Language | What are drugs that associate with Osteoporosis? |  |  |
| 3*<br>Query<br>set 3 | Parameter | Answer entity type | Question entity type | Question entity name |
|  | Input value | Drug | Gene | ESR1 |
|  | Natural Language | What are drugs that associate with gene ESR1 |  |  |

### G. Query result

Based on the parameters entered by the user, BEE displays three kinds of information. The first shows the network visualization, the second is the query result table after applying the query search operators, and the third is the result of each separate query set.

The query result table(Figure 4) displays the calculation results processed by the selected operation after the query entities entered by the user are converted to answer entities. The Co-occurrences column(Figure4-3) shows the number of genes shared with the answer entity that matched the transformation of the question entity.

**Fig. 2.**
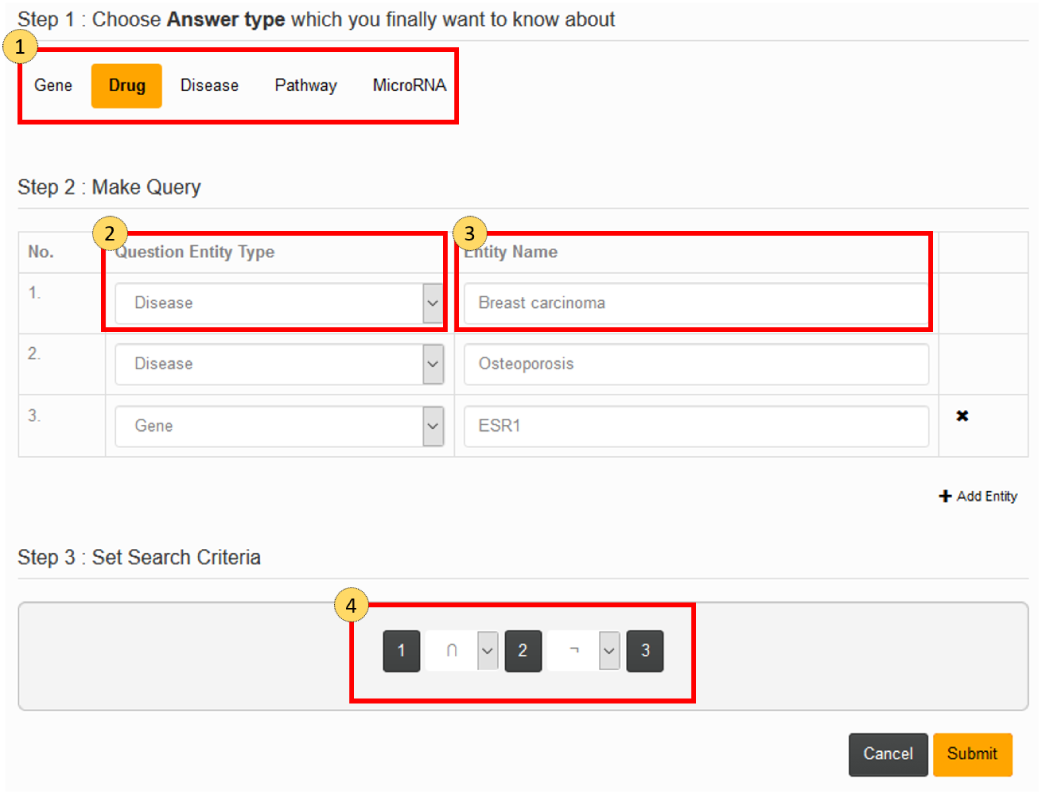
Query interface of BEE.

**Fig. 3.**
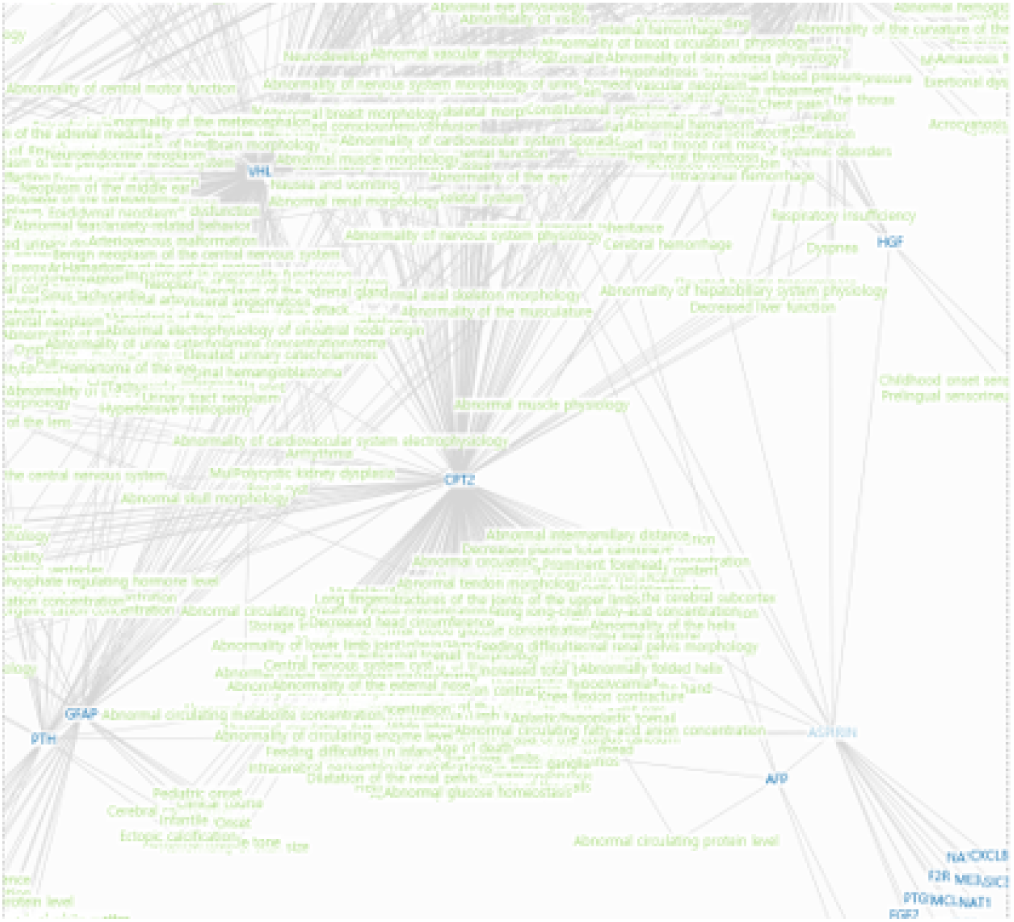
Query result : network visualization.

**Fig. 4.**
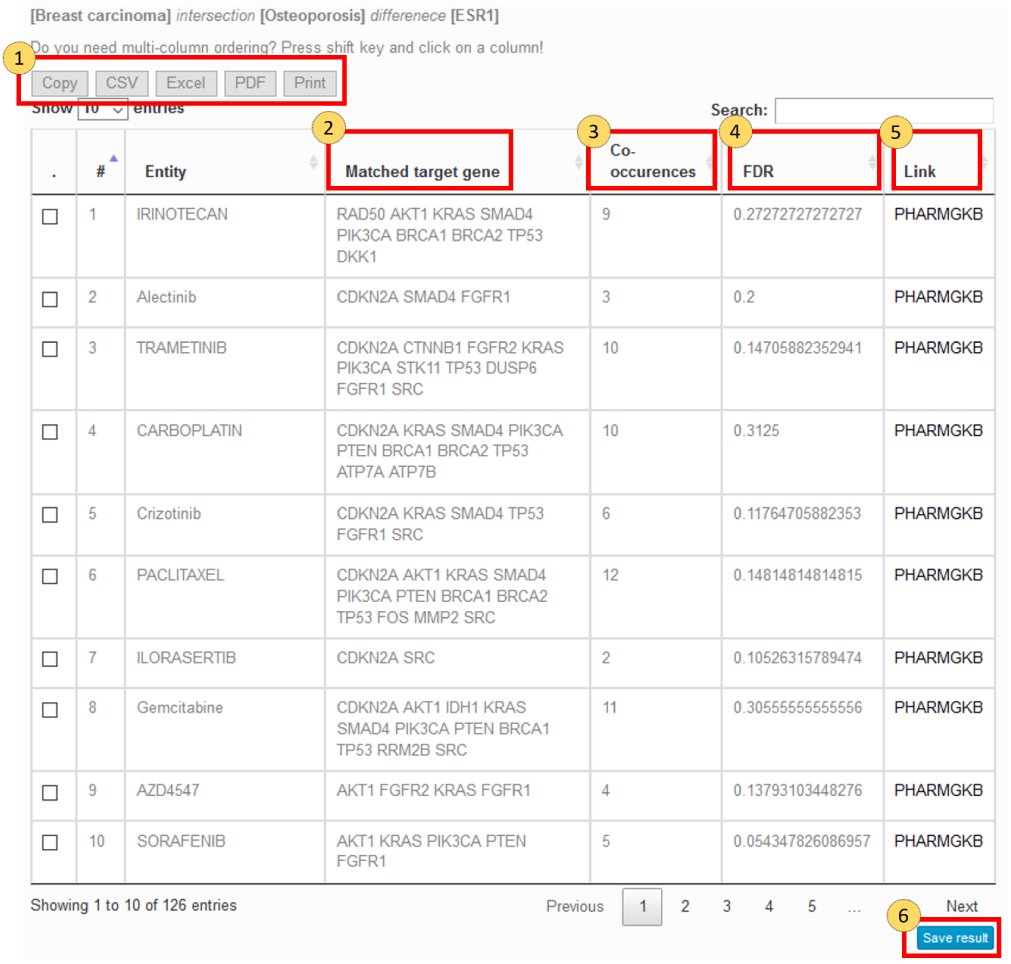
Query result table.

For instance, the input queries are *d*_1_, *p*_1_ and the operator is sum, as in Figure 5. The user can get the result set *ext*(*T*_*S*_, *rgen*(*gen*({*d*_1_}))) ∪ *ext*(*T*_*S*_, *rgen*(*gen*({*p*_1_}))) = {*s*_1_, *s*_2_, *s*_3_, *s*_4_, *s*_5_, *s*_7_}. The co-occurrence value of each answer entity is |(*gen*({*d*_1_}) ∪ *gen*({*p*_1_})) ∩ *gen*({*s*_1_})|, FDR value is *co* − *occurrenece/*|*gen*({*s*_1_})|.

**Fig. 5.**
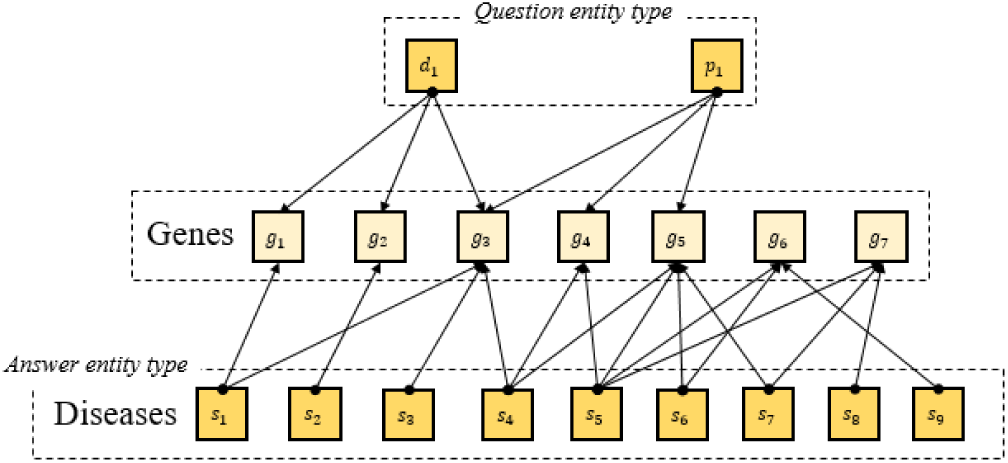
Query process.

The Link column (Figure 4-5) links to a website that provides details about the answer entity type. Services provided are BioPortal(8), GeneCards(9), PHARMGKB(10), Reactome, MirBase, Disgenet(11). In addition, the results can be copied to the clipboard, CSV, Excel, PDF, Web print. In addition, BEE provides feature which save search results for logged-in users (Figure 4-6). From the output, users can save selected entities and calculation results along with the description. When creating another associated query, users can create a chain query by loading and adding previous saved results. This leads to a different form of query from a previously typed question, which helps the form of the query proceed in various directions.

## Discussion and future directions

The current version of BEE has some limitations. First, the pathway uses not only a gene linkage but also various factors such as inclusion relations between pathways and … changes due to chemical interactions of proteins. In addition, data representation differs according to the viewpoint of the researcher. Currently BEE only considers genes that interact with the path with extreme abstraction because of the lack of a standard schema for representing data. In the next version, various schemas such as KEGG, Reactome, GO, etc. are considered, and data structures reflecting model reification will be applied. Not only pathway but also the drug and disease data remains unsatisfactory. Drugs and diseases have different names and levels in a country or institution, but the current version of BEE does not reflect mappings for various terms. Like the aforementioned pathway data, this also needs to be fixed in the next version. In addition, by providing a filter according to data source in the configuration of a question entity, we intend to provide a function for deriving query results from specific data sources.

Although there are limitations in the current version, BEE was developed to provide a simple and intuitive interface to easily answer complex queries about the association of polymorphic entities. In addition, it provides a visual representation of the network of query results and improves usability by supporting the output of search results in various data formats. In addition to simply retrieving the results of the query, various derivative results can be obtained by supporting chain-query which can be stored and loaded with other queries. As a result, BEE is expected to be used not only in the search for relationships among entities, but also in various fields such as drug repositioning.

## Notes

http://bike-bee.snu.ac.kr

